# Ecosystem tipping points in an evolving world

**DOI:** 10.1101/447227

**Authors:** Vasilis Dakos, Blake Matthews, Andrew Hendry, Jonathan Levine, Nicolas Loeuille, Jon Norberg, Patrik Nosil, Marten Scheffer, Luc De Meester

## Abstract

There is growing concern over tipping points arising in ecosystems due to the crossing of environmental thresholds. Tipping points lead to strong and possibly irreversible shifts between alternative ecosystem states incurring high societal costs. Traits are central to the feedbacks that maintain alternative ecosystem states, as they govern the responses of populations to environmental change that could stabilize or destabilize ecosystem states. However, we know little about how evolutionary changes in trait distributions over time affect the occurrence of tipping points, and even less about how big scale ecological shifts reciprocally interact with trait dynamics. We argue that interactions between ecological and evolutionary processes should be taken into account for understanding the balance of feedbacks governing tipping points in nature.

## Tipping points in an evolving world

Tipping points mark the abrupt shift between contrasting ecosystem states (broadly termed regime shifts) when environmental conditions cross specific thresholds (Box 1). Prominent examples are the shift of shallow lakes from a clear to a turbid water state (Scheffer et al. 1993), or the collapse of vegetation to a desert state in drylands (Reynolds et al. 2007). Societal stakes associated with tipping points in natural ecosystems can be high and there is great emphasis on the mechanisms that trigger them (Oliver et al. 2015) and the possible ways to detect and avoid them (Scheffer et al. 2009). Currently, however, tipping point theory lacks an evolutionary perspective, and this might limit our understanding of the occurrence, timing, and abruptness of shifts between states (Figure 1). Here we argue that both trait variation and evolution are important for understanding ecosystem dynamics in the vicinity of tipping points.

**Figure 1.**
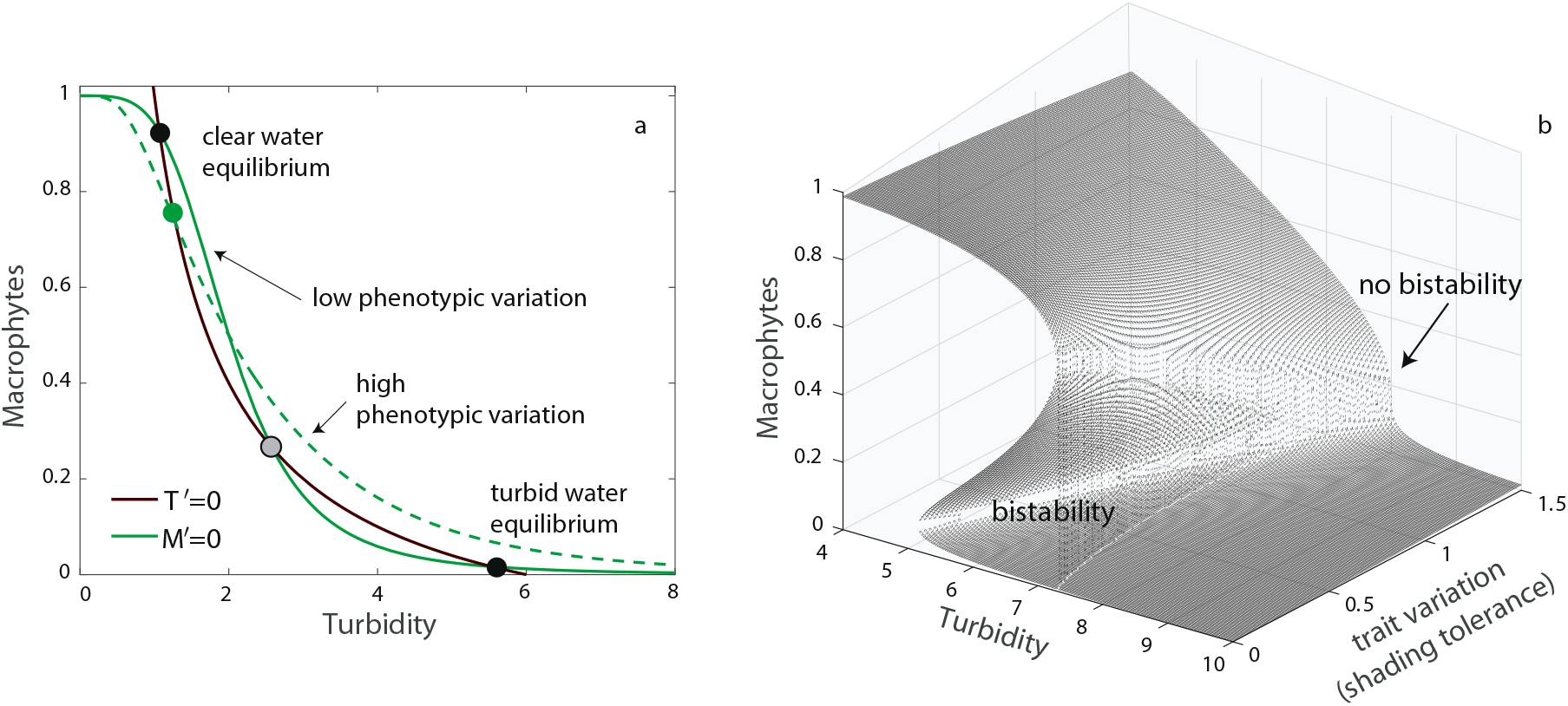
Variation in a response trait (eg macrophyte shading tolerance) affects tipping points of shallow lake shifting to a eutrophic turbid state. a) The intersections of macrophyte and turbidity responses (*M*’= 0, *T*’= 0 nullclines) mark the equilibria of the system for two levels of trait variation in the shading tolerance of macrophytes. In the absence of variation (*σ*^2^ = 0) there are two alternative equilibria (clear water and turbid water state at the crossing of solid green and brown lines). In the presence of variation (*σ*^2^ = 0.75), there is only a single equilibrium of clear water state with no tipping points (at the crossing of dashed green and solid brown lines). b) Changing the level of trait variation in the response trait (eg shading tolerance) will affect the response of a shallow lake to environmental stress (turbidity). Under increasing trait variation hysteresis decreases, bistability disappears, and the tipping point turns into a gradual and non-catastrophic response. Although not captured explicitly by this simple model, the effect of trait variation on the ecosystem response could act through the existence of resistant individuals (or subpopulations of macrophytes), but also on its potential to facilitate trait change. Model details and parameters can be found in the Supplementary Information.

Developing a trait-based evolutionary perspective about tipping points in ecosystems is warranted by the growing evidence that changes in standing levels of trait variation and contemporary trait evolution are important drivers of ecological dynamics (eg (Saccheri and Hanski 2006; Kinnison and Hairston 2007)), influencing population dynamics (Yoshida et al. 2003), shaping the structure of species interactions and composition at the community (Pantel et al. 2015), or at metacommunity level (Farkas et al. 2013). Such ecological effects of evolution also extend to ecosystem functioning (Norberg et al. 2001; Matthews et al. 2011; Hendry 2017), by modifying material fluxes (De Mazancourt et al. 1998), primary production (Gravel et al. 2011), nutrient recycling (Loeuille et al. 2002), and decomposition (Boudsocq et al. 2012). Thus, it is reasonable to expect that trait distributions could be important for ecological tipping points by affecting response diversity in an ecosystem; that is variation in the sensitivity among species, populations, or individuals to environmental stress (Elmqvist et al. 2003). This sensitivity underlies the response capacity of communities to stress (Vellend and Geber 2005), such that trait change could affect the resilience of entire ecosystems to stress (Mori et al. 2013) and their probability of tipping to a different state.

Ecosystem resilience can be affected by variation in traits (Norberg et al. 2001; Matthews et al. 2011) underlying the performance and fitness of organisms in a given environmental state (i.e. response traits), or those causing direct or indirect effects on the environmental state (i.e. effect traits) (Table 1). The distribution of such response and effect traits can vary due to phenotypic plasticity or evolutionary trait change, and distinguishing between these mechanisms can be important for understanding the temporal dynamics of trait change in general (Cortez 2011), and of tipping points in particular. Phenotypic plasticity, where genotypes exhibit different phenotypes in different environments, is a relevant source of trait variation, particularly when the phenotypic changes relate to the capacity of organisms to respond to stress. However evolutionary responses to stress depend on heritable trait variation in a population (Hansen et al. 2012), which can originate from novel variants due to mutation (Nei 2007), recombination (Ortiz-Barrientos et al. 2016), or gene flow among populations and species (Seehausen 2004). Below, we do not *a priori* distinguish between the genetic versus plastic sources of trait distributions (although we comment on their differences), but focus on how trait variation and trait change over time can influence ecosystem tipping points in a generic way. We do this using a graphical approach where we illustrate how trait changes might modify the collapse and recovery trajectories of ecosystems along an environmental gradient.

**Table 1.**
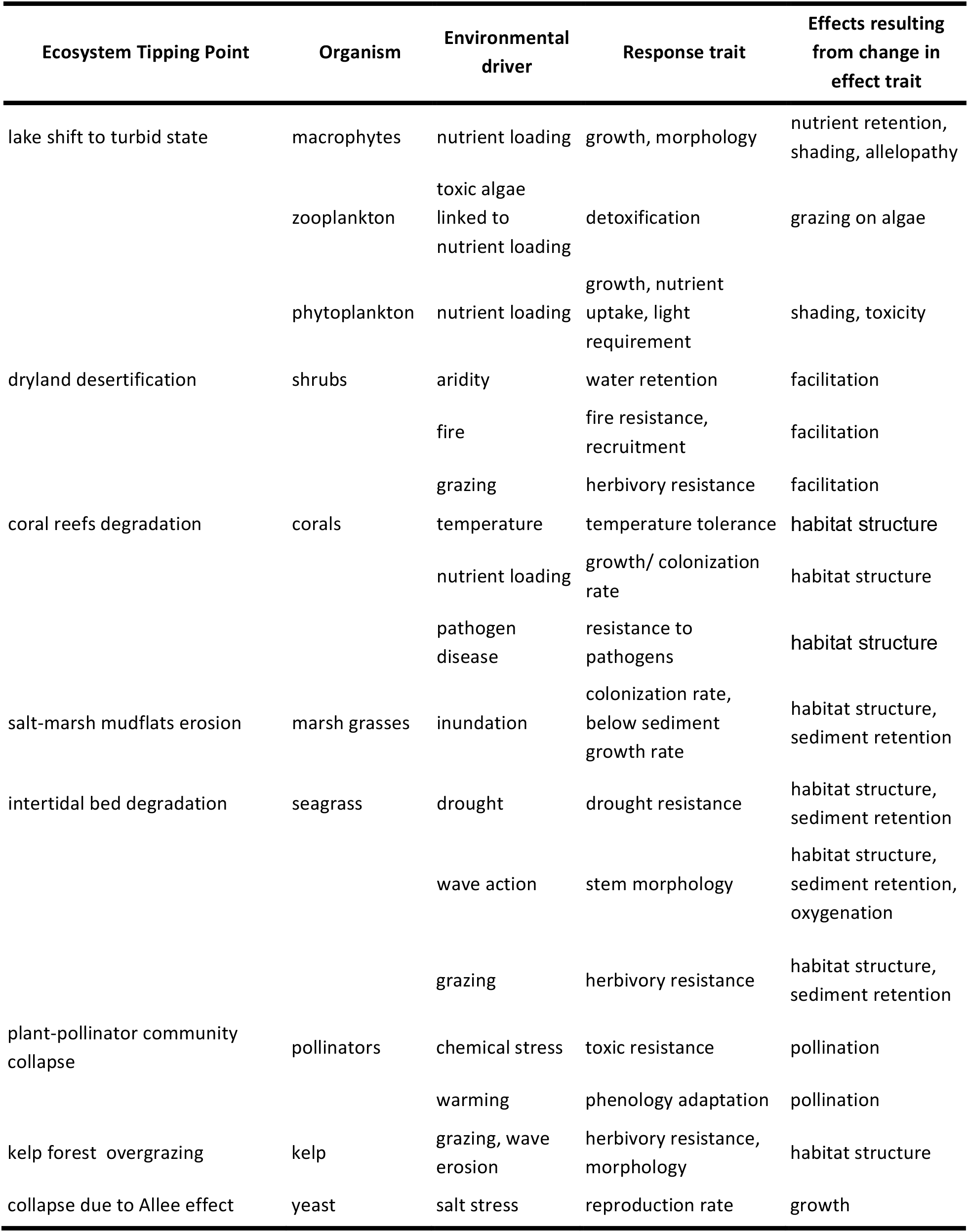
Examples of ecosystem tipping points summarizing the organisms involved and the potential response and effect traits of these organisms. If these traits can experience phenotypic changes, they may affect the tipping point responses in any of the ways presented in the text. Response traits are defined as traits that respond to the environmental stressor(s) that can invoke a tipping point. Effect traits are defined as traits that may influence an ecosystem function that is linked to a tipping point. In the table we refer to the effect of such traits rather than the traits themselves.

### Trait variation could affect the probability of tipping points

Differences in the amount of trait variation in populations could affect their response capacity to stress. In general, we predict that high trait variation may decrease the probability of tipping points turning ecosystem responses to non-catastrophic. A decrease in the probability of tipping events occurs because standing trait variation allows for portfolio effects that introduce strong heterogeneity in population processes, interactions, and responses (Bolnick et al. 2011) buffering population dynamics (Schindler et al. 2010). Such heterogeneity can be enhanced by Jensen’s inequality (Bolnick et al. 2011), where variation around the mean of a trait can affect the response of an ecological interaction or an ecological process in function of the nonlinear relationship between the trait and its effect (Ruel and Ayres 1999). This effect can be clearly illustrated in a toy model describing shifts in the case of shallow lakes (Figure I in Box 1). Here, changing the amount of variation in the macrophytes’ response trait to turbidity can increase or decrease the probability of a tipping point response. Under high levels of variation the transition from the clear to the turbid water state can even become non-catastrophic with no alternative states (Figure 1).

### Trait change could delay a tipping point

As introduced in the previous paragraph, trait variation simply means that some resistant phenotypes are present. However, trait variation could also facilitate trait changes. On top of that, trait changes might be fueled by *denovo* mutation and phenotypic plasticity. In ecosystems where stress gradients bring them closer to tipping points, trait changes could potentially delay tipping to the alternative state (Figure 2a). This resonates with the idea of evolutionary rescue (Gomulkiewicz and Holt 1995), the difference being that there is no rescue, but rather only a delay in the collapse of the system by shifting the threshold at which the collapse occurs at a higher stress level (Figure 2b). For instance, in the case of a shallow lake turning turbid due to eutrophication (Box 1), aquatic macrophytes might delay the transition to a higher threshold level of nutrients because of contemporary changes in traits that convey tolerance to shading (Table 1).

**Figure 2.**
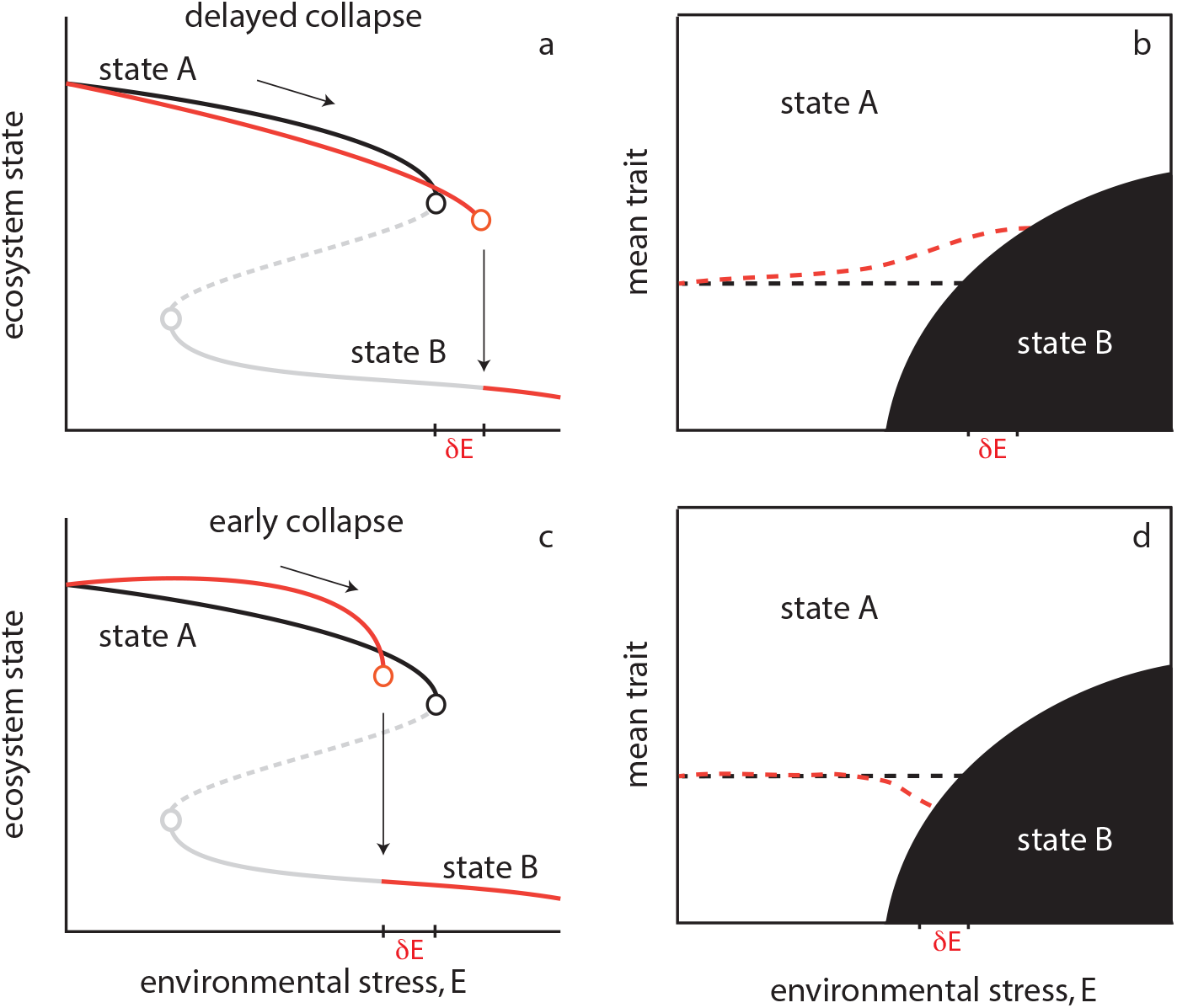
Hypothetical alterations of trajectories of ecosystem collapse (left panels, red solid lines) as a consequence of trait change (right panels, red dotted lines). (a, b) Contemporary adaptive mean trait change delays the threshold at which the tipping point occurs (δE), potentially associated with a cost that decreases the equilibrium ecosystem state. (c, d) Adaptive mean trait changes might in the short term increase the equilibrium ecosystem state while at the same time also induce an early collapse. [(a, c) Black and gray lines represent the two alternative states of the reference model with no phenotypic change, dotted lines mark the unstable boundary between the two states, dots denote tipping points. (b, d) Dotted black line is the reference scenario with no trait change]

### Trait change could lead earlier to a tipping point

Trait change may not always buffer populations from environmental changes, but could also contribute to an increased risk of ecosystem collapse (Figure 2c, d). For example, environmental stress could impose directional selection on a trait in a given species or group of species that brings the system closer to tipping to an alternative ecological state (Dieckmann and Ferriere 2004; Rankin and Lopez-Sepulcre 2005). This is similar to evolutionary collapses or evolutionary suicide as defined in evolutionary biology (Ferriere and Legendre 2013), but here the collapse occurs at the scale of a whole ecosystem. Empirical examples of trait evolution leading to population collapse come mostly from fish populations under harvesting (Rankin and Lopez-Sepulcre 2005; Walsh et al. 2006). For example, it has been shown how fishing pressure has led to the early maturation of Atlantic cod populations (Olsen et al. 2004) that is associated with lower reproductive output and irregular recruitment dynamics that could have increased the chance of stochastic extinction and the cod collapse in the 1990s. Evolutionary suicide might lead to an ecosystem-level collapse in the case of drylands (Kéfi et al. 2008), where under increased aridity adaptive evolution can favor local facilitation among neighboring plants for resisting higher aridity. Whether evolution leads to a buffering effect depends on the seed dispersal strategy of the dominant vegetation type. In systems characterized by long-distance dispersal, evolution may actually enhance the collapse of the vegetation to a desert state due to the invasion of non-facilitating mutants. In our shallow lake example, macrophytes at intermediate turbidities might respond by growing longer stems with fewer leaves in order to reach well-lit surface waters and avoid shading. If this, however, results in less photosynthetic activity and less capacity to remove nutrients from the water column, it might reduce the capacity to outgrow the algae and maintain a clear water state.

### Trait change could affect the path of recovery

Changes in trait distributions over time may also affect the recovery trajectory of an ecosystem back to its previous state and the range of hysteresis, i.e. the lag in the threshold of the environmental driver at which recovery to the pre-collapsed state occurs (see Box 1 and Box 4 (Glossary)). The most obvious example is the case where trait change delays a tipping point (Figure 3). In many cases, this delay will not necessarily result in an equally early recovery, which implies that hysteresis in the system will increase. This example illustrates that tipping points and hysteresis are the flip side of mechanisms buffering the stable states: if evolution or phenotypic plasticity buffers the system against environmental change, this can not only delay reaching a tipping point but it may also result in stronger hysteresis.

**Figure 3.**
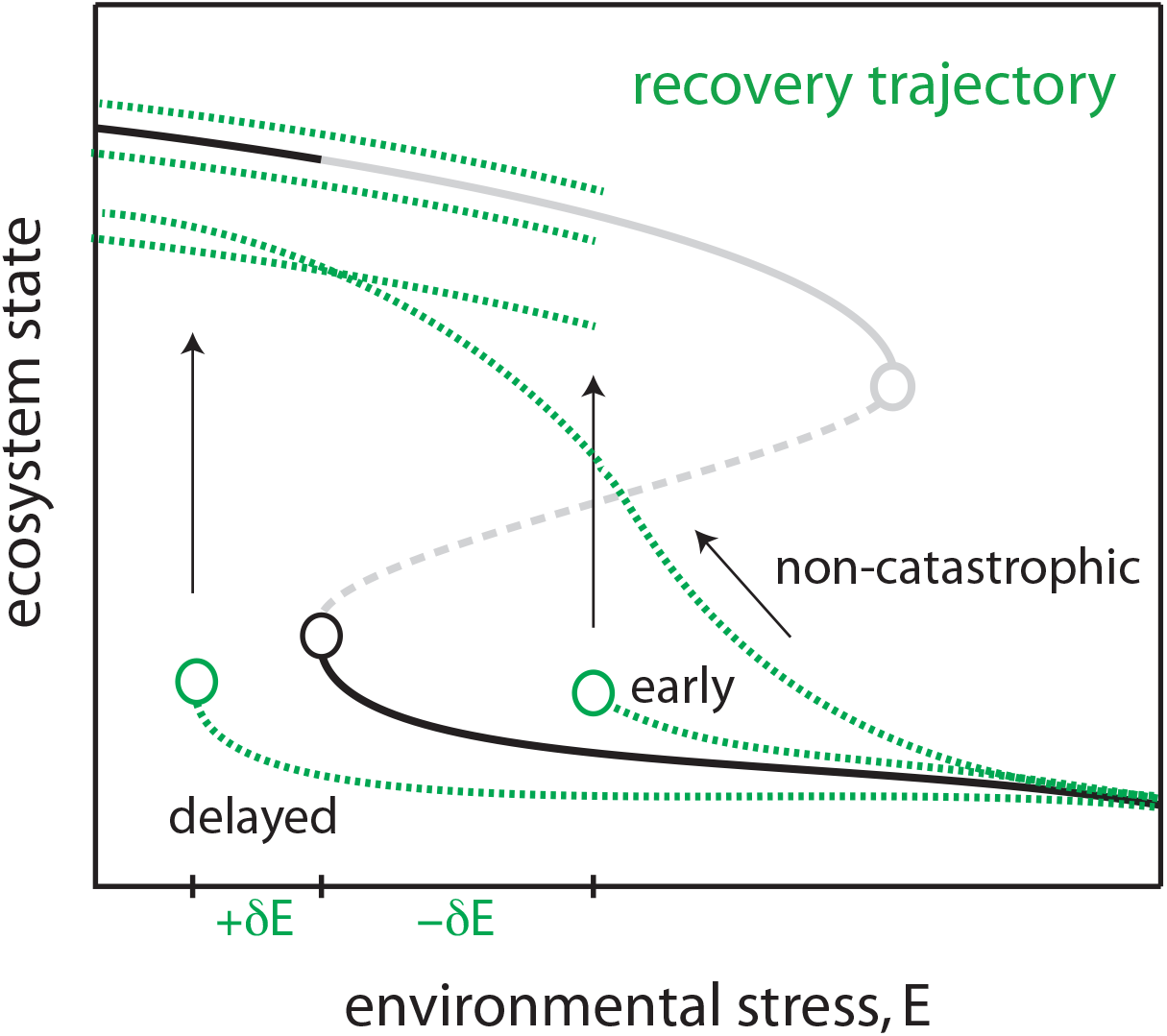
Potential consequences of trait change on the recovery trajectories of an ecosystem after collapse (green dashed lines). Recovery may be delayed or occur earlier affecting the range of hysteresis and the ease of recovery. In both cases, it is unclear whether the ecosystem shifts back to exactly the same state as before the collapse. It may be possible that the collapse has allowed the emergent of a different (new) phenotype that could even turn the recovery path non-catastrophic. [Solid lines represent the two alternative states of the reference model with no phenotypic change, dotted lines mark the unstable boundary between the two states, dots denote tipping points.]

Another possibility is that evolutionary processes in the deteriorated state might cause the collapsed species to lose the genetic variation necessary for recovery to, and high fitness in, the alternate state (Murray et al. 2017). In a laboratory experiment, (Walsh et al. 2006) found that overharvested fish populations failed to recover even after reducing fishing pressure due to genetic changes in life history traits. This may result in a delay in recovery, or no recovery at all. The opposite scenario is also possible. Trait changes may accelerate recovery and reduce hysteresis (Figure 3). This may happen if, after the collapse, a highly adaptive phenotype is selected for facilitating recovery only at a small reduction of stress. For example, after the collapse of a phytoplankton population due to light stress in the laboratory, recovery took place earlier than expected due to a (probably plastic) adaptive photo-acclimation response (Faassen et al. 2015). If after the collapse a different phenotype is selected for, or if there is recovery of the lost phenotypic variation (e.g. due to immigration), it may even be possible that the recovery pattern becomes non-catastrophic.

In all cases highlighted in the previous paragraphs, it is uncertain whether the ecosystem will actually recover to the exact same state as before the collapse (Figure 3). The degree to which complete recovery happens might probably depend on the trait that changes. It is one of the outstanding questions whether trait changes that impact the probability of tipping also impact recovery trajectories (Box 3).

### Phenotypic plasticity, evolution and tipping points

There are more possibilities for the collapse and recovery paths of the ecosystem state than the ones we highlighted here. All will depend on the mechanisms of phenotypic change and it requires both theoretical and empirical work to understand the most probable outcomes on tipping point responses that would result either from evolution, from phenotypic plasticity, or from their combined effect, including even the evolution of phenotypic plasticity. One reason why the distinction between phenotypic plasticity and evolutionary trait change is important is that the rates at which these processes operate tend to differ, with phenotypic plasticity being generally faster than evolutionary change. Conversely, phenotypic plasticity is often limited in amplitude, and evolutionary trait change might extend the range to which tipping points and hysteresis can be impacted. Importantly, trait change due to evolution also has an intrinsic impact on the population genetic structure that entails a legacy that may impact recovery (e.g. case of genetic erosion or a trait change that is adaptive in one stable state but maladaptive in the other state), whereas trait change mediated by phenotypic plasticity may impact tipping points without a legacy effect if the trait change is reversible.

## Testing the effects of phenotypic change on tipping point responses

### Integrating evolutionary dynamics in models of ecological tipping points

Coupling models on evolutionary dynamics with models of ecological bistability can offer a better understanding about when genetic trait change can affect tipping point responses. The adaptive dynamics framework - that assumes limited mutation and the separation of ecological and evolutionary timescales - has been used to study how evolution may incur evolutionary collapse and suicide (Dieckmann and Ferriere 2004). Under rapid environmental change, a quantitative genetics approach (Abrams 2005) is useful for studying how contemporary genetic trait change may lead to evolutionary rescue. Both modelling frameworks can be adapted for studying how trait changes might affect well-understood models with ecological tipping points under changing environmental conditions. For instance, we could relax the assumption on the separation of timescales and the assumption of weak selection of each framework, respectively, and apply them to models with tipping points. Or one could develop hybrid models that can account simultaneously for selection gradients, while also accounting for genetic drift and demographic stochasticity that dominate the recovery trajectory of the collapsed state. We can then combine these models with recently developed methods that measure the relative impact of evolutionary vs ecological dynamics on stability (Patel et al. 2016) to understand when and how evolutionary dynamics can affect the probability of tipping responses.

Such modelling approaches can help to (i) compare how different mechanisms of trait change (genetic vs plastic) could affect tipping point responses, (ii) identify the conditions (e.g. rate and pattern of environmental stress, rate of trait evolution, costs and trade-offs) under which trait evolution will modify collapse and recovery trajectories, or even (iii) test when trait change itself could be so abrupt (due to disruptive selection) that it could cause ecosystem tipping points. In that way we could develop novel ways for detecting tipping points based on changes in ecological and trait dynamics (Box 2), and suggest new designs for experimental testing.

### Adding evolutionary contrasts to experimental tests of ecological tipping points

There are two common approaches for experimentally testing tipping point theory. The first approach starts by establishing two alternative states of the system on either side of a tipping point, and then testing how the system responds to pulse perturbations of a state variable. For example, if there is evidence for a positive feedback (Box 1) in two states with a different dominant species in each community, then the outcome of species dominance might strongly depend on the initial density of species (i.e. priority effects) (Fukami and Morin 2003). The second approach starts with the system in one state, and then applies a press perturbation of an environmental condition (e.g. increasing productivity, increasing mortality) to observe when the system transitions to a new state (Dai et al. 2012; Veraart et al. 2012). To test for hysteresis in the system, the environmental condition can then be reversed while tracking system recovery to the initial state (Faassen et al. 2015).

Independently manipulating evolutionary and ecological components of a system can provide new insights into how the dynamics of trait change can affect tipping points. Several experiments have been designed to study the interplay between ecological and evolutionary dynamics (Farkas et al. 2013; Pantel et al. 2015; Williams et al. 2016), and these could be usefully co-opted to experimentally test predictions from tipping point theory. In an experiment with freshwater cyanobacteria, light level was manipulated to test for hysteresis associated with transitions between a high and low biomass state (Faassen et al. 2015). Contrary to predictions from an ecological model, the population recovered to a higher light stress faster than expected. In the experiment, the recovering cells had lower pigment concentrations, possibly reflecting adaptation to high irradiance conditions at a cost of photosynthetic efficiency at lower light irradiance. This suggests that the presence of trait variation in the population influenced the nature of the transition between the two states. A useful experimental test of this idea would be to manipulate standing levels of genetic variation in the stressed population and measure if tipping points change. Adding such evolutionary contrasts to ecological experiments would be a fruitful way to test how both trait variation and evolution may affect tipping points. In experimental systems it is possible to isolate the effects of density (ecological effects) from the effects of heritable trait change (evolutionary effects). Specifically, one might be able to differentiate between purely ecological effects, direct evolutionary effects linked to changes in functional effect traits, and density-mediated indirect evolutionary effects linked to changes in functional response traits (Patel et al. 2016).

## Closing the loop: eco-evolutionary feedbacks and tipping point responses

Reciprocal interactions between ecological and evolutionary dynamics is an old idea (e.g. (David 1968; Levins 1968)) that is increasingly being tested across a range of systems and study questions (e.g. (Fussmann et al. 2007; Hendry 2017)). Here, we focused on the potential implications that heritable trait changes can have for ecological tipping points. The next step is to understand how reciprocal feedbacks between ecological tipping points and evolutionary dynamics might radically alter not only the dynamics of ecosystems close to tipping but also the evolution of populations and communities of these ecosystems. Tipping points between contrasting ecosystem states create different selection regimes that can shape the evolution of focal species (like keystone, or ecosystem engineers species) and in their turn the dynamics of the ecosystem state they belong to (Matthews et al. 2015). One possibility is that such selection regimes will be asymmetric, leading to evolutionary reversals, for example in body sizes in grazed populations (Dercole et al. 2002), or could maintain the recurrence of harmful algal blooms in lakes (Driscoll et al. 2016).

It remains an outstanding challenge to test these ideas along with several new questions (Box 3). Most theoretical work on eco-evolutionary dynamics has been experimentally corroborated in laboratory experiments with short generation organisms (Yoshida et al. 2003). Similarly, ecological tipping points have been mostly studied in experimental microcosms at the population level with single species (Dai et al. 2012; Veraart et al. 2012). Ecosystem scale tipping points are harder to experimentally test (but see (Carpenter et al. 2011)) and simultaneous information on trait variation of the organisms involved is rarely available. Yet, ecosystem collapses have evolutionary consequences that may trap an ecosystem in an undesired state making recovery difficult. Thus, sustaining trait variation may be important not only for preventing collapse, but also for improving the success of ecological restoration. Despite the challenging task, the evolutionary perspective we advocate can improve our understanding and management of ecosystems under stress.

## Acknowledgements

VD and BM are grateful to Eawag and the Adaptation to a Changing Environment Program at ETH Zurich for financing a workshop on eco-evolutionary dynamics of tipping points help in Kastanienbaum in 2016. BM acknowledges a SNF 31003A_175614 grant.

## Competing interests

We declare no competing interests.

## Author contributions

VD and BM designed research and wrote the paper with contributions from all authors.

### Box 1 What is a tipping point?

Tipping points mark the shift between contrasting system states that occur when external conditions reach thresholds that trigger an accelerating transition to a contrasting new state (Nes et al. 2016). Mathematically, these transitions correspond to saddle-node or fold bifurcation points (Strogatz 1994). They are also called catastrophic because they mark an unexpected and radical change in the equilibrium state of a system. Tipping points can occur at population level (e.g. due to Allee effects (Dai et al. 2012)) and community level (e.g. due to trophic cascades (Kitchell and Carpenter 1993)), but it is at the ecosystem scale that tipping points are most prominently studied because they can incur long-term disruption to vital ecosystem services (Barnosky et al. 2012). For example, clear lakes turn turbid dominated by algal blooms (Scheffer et al. 1993), coral reefs get overgrown by macroalgae (Knowlton 1992), fisheries collapse due to overexploitation (Beddington and May 1977), and tropical forests shift to savanna-type ecosystems under high fire intensity (Staver et al. 2011).

Tipping points are typically observed in systems where strong positive feedbacks drive the establishment of alternative stable states (Nes et al. 2016). In the case of shallow lakes, dominance of aquatic macrophytes prevents the growth of algae by removing nutrients (phosphorus) from the water column that leads to the establishment of a stable clear water state (Fig I). When phosphorus loading exceeds a critical threshold macrophytes cannot successfully retain phosphorus, algae start to grow and lake turbidity increases. Rising turbidity kicks a vicious cycle: it hinders the growth of macrophytes but facilitates algae concentration in a self-enforced positive feedback loop (less macrophytes => more algae => more turbidity => less macrophytes and so on) that leads to the collapse of macrophytes and the establishment of a contrasting turbid lake state. The same positive feedback loop can lead to the recovery of macrophytes, but this time at a lower critical level of phosphorus loading, where algae growth is limited to such an extent that turbidity decreases sufficiently for macrophyte to grow again, capture the phosphorus and reinforce a positive feedback loop leading back to the clear water state. Between these two tipping points, the system is bistable meaning that it can be found in one of the two alternative stable states. This difference in conditions that mark the forward and backward shift is called hysteresis. The stronger the hysteresis, the more difficult it is to recover an ecosystem back to its previous state.

**Figure I.**
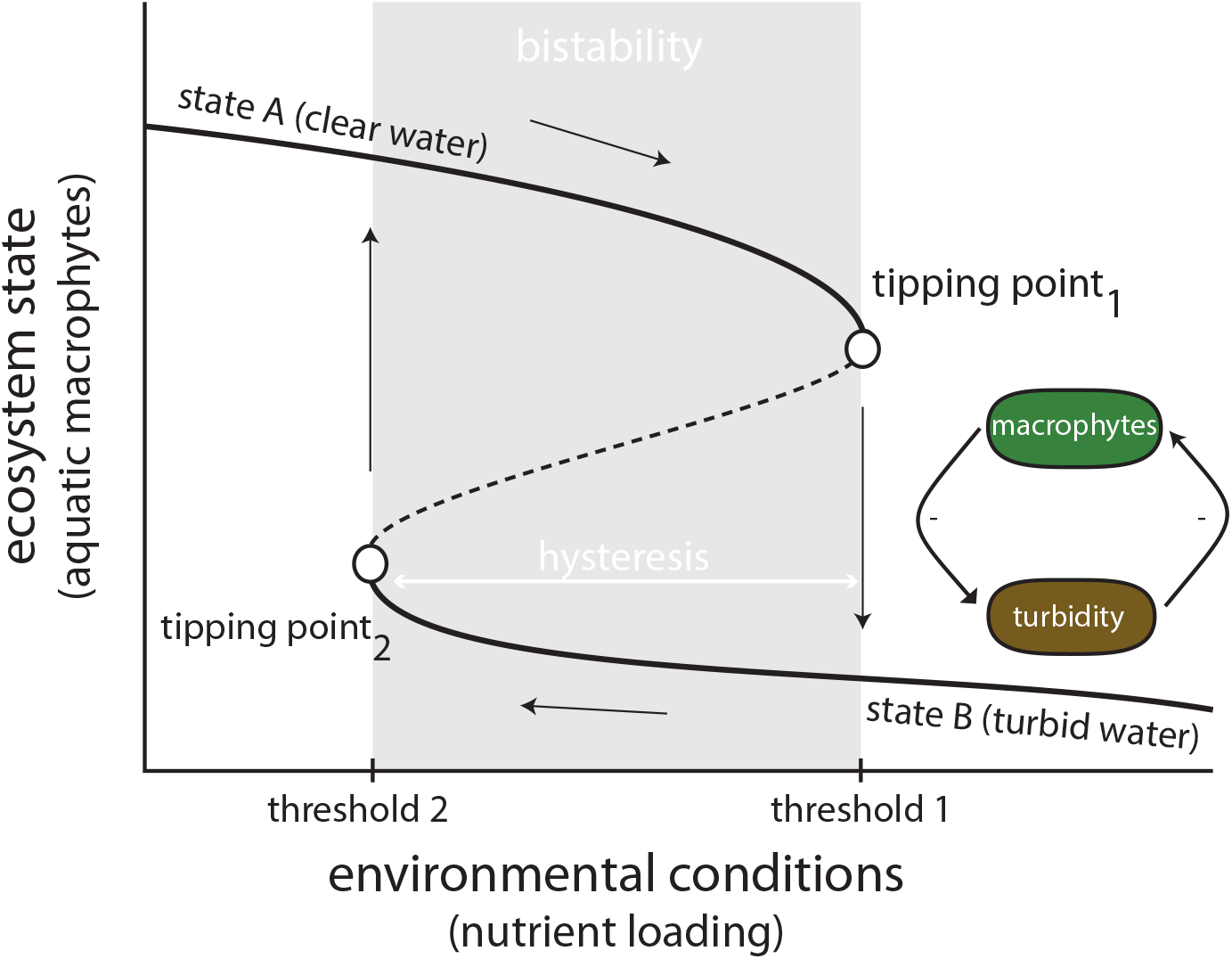
Tipping points mark discontinuous changes in the state of an ecosystem. Starting from the upper branch, the ecosystem follows the stable equilibrium line until conditions cross threshold 1 at which the upper stable equilibrium disappears (tipping pointi) and the ecosystem state drops abruptly to the lower (alternative) stable state. In our example of the turbid and clear-water states of shallow lakes, reducing nutrient conditions - but to a much lower level - leads to the restoration of the previous state at the crossing of threshold 2 (tipping point_2_). The difference in the thresholds between the forward and backward tipping points marks the hysteresis in the system. For this range of conditions the ecosystem can be found in either of the two alternative stable states (bistability). Along the pathways depicted here, no change in the traits of the organisms stabilizing the clear-water (macrophytes) or turbid (algae) state is assumed. [Black lines represent the stable equilibria. Dotted line represents the border between the basins of attraction of the two alternative stable states.]

### Box 2 Detecting tipping points based on ecosystem-state and trait changes

Ecological tipping points are difficult to detect. However, theory suggests that subtle changes in the dynamics of an ecosystem state can provide early-warning information on the underlying stability and risk of a tipping response (Scheffer et al. 2009). This risk is typically quantified by indicators of resilience based on critical-slowing-down (Dakos et al. 2015), and include an increase in recovery time back to equilibrium after a perturbation, a rise in variance as the state of the ecosystem fluctuates more widely around its equilibrium, and an increase in autocorrelation because the state of the ecosystem resembles more and more its previous state close to a tipping point. These indicators have been empirically tested in laboratory experiments (Dai et al. 2012; Veraart et al. 2012) and in the field (Carpenter et al. 2011; van Belzen et al. 2017) focusing on ecosystem state, and neglecting any trait changes. Accounting for trait change creates new challenges but also opportunities in the detection of tipping points (Figure II). It is unclear whether trait change can affect the performance of resilience indicators, or whether indicators based on both ecosystem state and trait state dynamics could complement each other to improve tipping point detection. Changes in traits have been suggested as a basis for predicting ecological responses (Enquist et al. 2015), and seeds of this idea can be found in the suggestion that variation in maturation schedules of cod could have been used to detect its collapse (Olsen et al. 2004). Recent work has shown that measuring changes in mean or variance in body size in combination with resilience indicators based on species abundance could improve warning of protists population collapse (Clements and Ozgul 2016). Nonetheless, slowing down indicators should be expected - at least based on ecological dynamics - in ecosystems at the edge of tipping points (Ferriere and Legendre 2013). Although changes in the dynamics of phenotypic adaptation will most likely be context-dependent, it remains to be tested whether they could be used as signals of potential impending transitions.

**Figure II.**
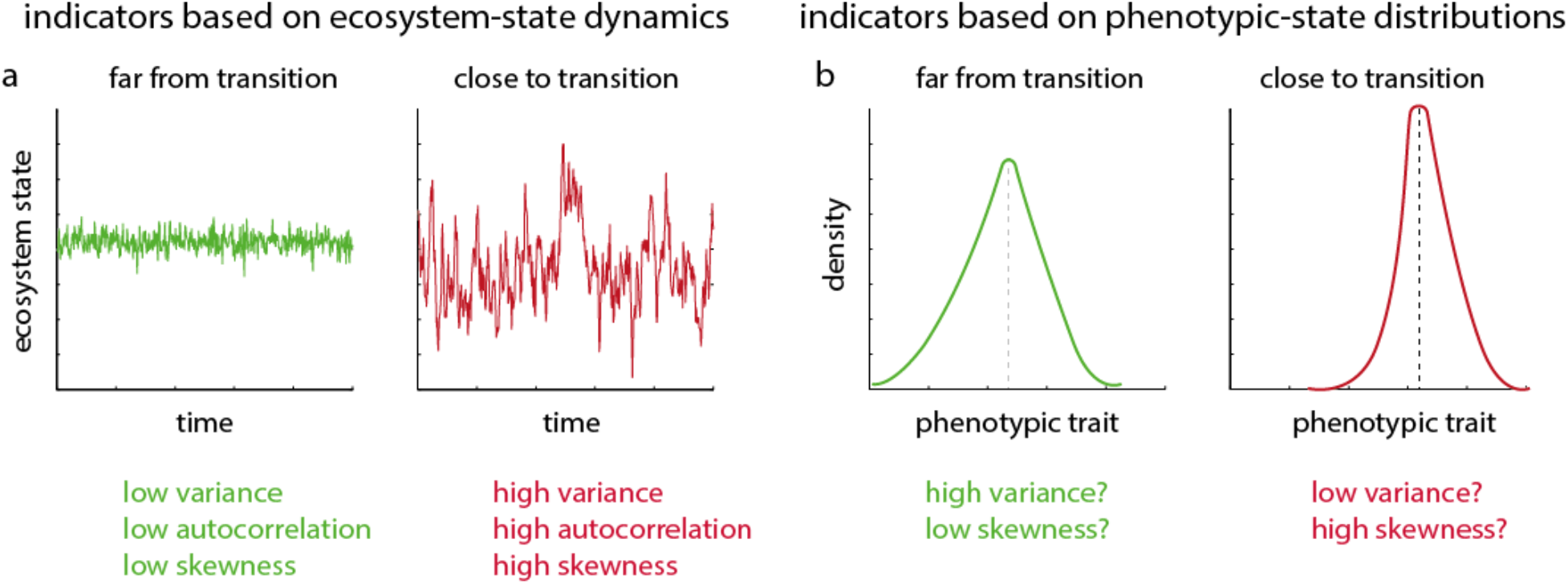
Deriving indicators of resilience for detecting the risk of tipping point responses from changes in ecosystems dynamics and trait distributions. Close to tipping points, ecological dynamics become slower in responding to disturbances. This slow recovery leads to a rise in variability and memory in the dynamics of the monitored ecosystem state that can be used as indicator of an increased risk of tipping (panel a). Alternatively to such ecosystem-state indicators, temporal changes or changes along a gradient in trait values of focal species might be also informative for quantifying the risk of approaching tipping points (panel b). Such trait based indicators may flag changes even before signals from ecosystem state flare up.

### Box 3 Critical questions for understanding tipping points from an evolutionary perspective

- Under which conditions (e.g. type and rate of environmental stress, type of response/effect trait, level of genetic variation, plasticity, spatial and temporal scales) does phenotypic change matter the most for ecological tipping points?
- In what ways do genetic vs plastic trait changes affect tipping point responses differently?
- Do reciprocal interactions between species (e.g. in a network) that influence trait changes (e.g. through coevolution) affect tipping point responses?
- Is there an intrinsic relationship between trait changes that impact collapse and recovery, and to what extent can trait changes that impact collapse can be reversed so as not to impact recovery?
- What type of eco-evolutionary feedbacks develop along the collapse and recovery trajectories of ecosystems with tipping points?
- Can ecological bistability lead to bistability in trait values (or vice versa)?
- Can we use changes in trait variation as signals of approaching tipping points?
- How can we experimentally study the effects of trait change in ecosystems with tipping points?
- Can we manage trait variation and evolution to reduce the risk of ecosystem tipping points?

### Box 4

**Glossary**
Alternative stable states: contrasting states that a system may converge to under the same external conditions
Bistability: the presence of two alternative stable states under the same conditions
Catastrophic bifurcation: a substantial change in the qualitative state of a system at a threshold in a parameter or condition
Contemporary (or rapid) evolution: evolutionary changes that occurs sufficiently rapid that it can have an impact on ecological dynamics at the same time-scale as other ecological factors
Eco-evolutionary dynamics: dynamics in which ecological processes influence evolutionary processes and evolutionary processes influence ecological processes
Effect trait: a measurable feature of an organism that underlies an organism’s direct effect on an ecosystem function
Genetic drift: changes in allele frequencies due to random sampling during reproduction
Hysteresis: the lack of reversibility after a catastrophic bifurcation, meaning that when conditions change in the opposite direction the system stays in the alternative state unless it reaches another bifurcation point (different than the one that caused the first shift)
Phenotypic plasticity: non-heritable changes in the phenotype of an organism
Response trait: a measurable feature of an organism that underlies an organism’s response to environmental change
Tipping point: the point where following a perturbation a self-propagated change can eventually cause a system to shift to a qualitatively different state
Trait variation: variability of any morphological, physiological, or behavioral feature
Trait evolution: genetic change in phenotype of a given trait

## Supplementary Information - Shallow lake eutrophication model

We used a minimal model that describes the dynamics of transition from a clear water state dominated by macrophytes to a turbid water state where macrophytes are practically absent^1^. Such transition occurs at a crossing of a fold bifurcation (tipping point) due to changes in nutrient loading (eutrophication). Below we explain how we analysed the model to highlight the presence of alternative states as function of environmental stress (Box 1), and the effects of standing phenotypic variation (Figure 1).

The model describes the interactions between macrophyte coverage and turbidity of a shallow lake with the following two ordinary differential equations:

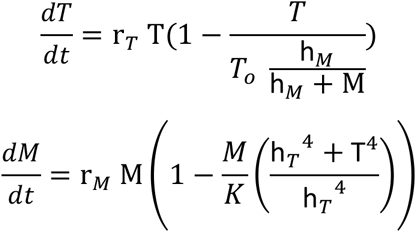

where macrophyte cover *M* grows logistically with rate *r_M_* (= 0.05) and carrying capacity *K* (= 1), while it is limited by turbidity following a nonlinear decreasing Hill function defined by the half-saturation *h_T_* (= 2) and exponent p (=4). Turbidity *T*grows with rate *r_T_* (= 0.1) depending on the level of background turbidity *T_o_* (= [2-8], used as proxy of nutrient loading acting as the environmental stress in our analysis (nutrient loading, Fig I Box I)). Turbidity is negatively affected by the level of macrophyte cover following an inverse Hill function with half-saturation *h_M_* (= 0.2).

Solving for steady state the nullclines of the system are:

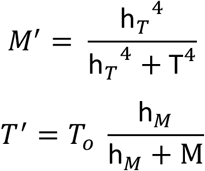

Their intersections mark the two alternative stable states (clear and turbid state) and the unstable saddle depending on the value of background turbidity *To* (Fig. 1a). We hypothesize that the half-saturation *h_T_* that affects the strength of nonlinear response of macrophytes to turbidity is defined by a response trait *z* (e.g. capacity to grow under low light conditionsshading). Different values of *z* will thus change the response of macrophytes to turbidity by changes in *h_T_* (Supplementary Figure 1a). We assumed that trait *z* follows a *beta* distribution (closed limits) that we can parameterize in order to define a given mean *μ* (=0) and variance *σ*^2^. We further assumed that the half-saturation *h_T_* depends on the trait *z* following *h_T_* = *h_To_e^cz^*, where *h_To_* is a background value (= 2) and *c* a factor (=0.5) (Supplementary Figure 1b).

Using this relationship and integrating for different limits of trait *z* and levels of variance of the *Beta* distribution, we can calculate the macrophyte equilibrium in the presence of standing phenotypic variation in z as:

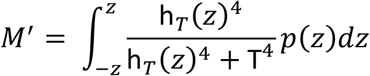

where *p*(*z*) is defined by the *Beta* distribution as explained above within a range of *z* (= [−2,2]). We repeat this for a range of turbidity *T* values (= [0-8]) to estimate the nullcline of macrophytes *M* for this range of turbidity *T*, and we find the new equilibria states from the cross sections with the turbidity nullcline (Fig. 1a).

We repeat this procedure to estimate all equilibria as a function of environmental conditions (*T_o_*) and for different levels of standing phenotypic variation (*σ*^2^) to construct the two dimensional bifurcation plot of Fig. 1b.

**Supplementary Figure 1.**
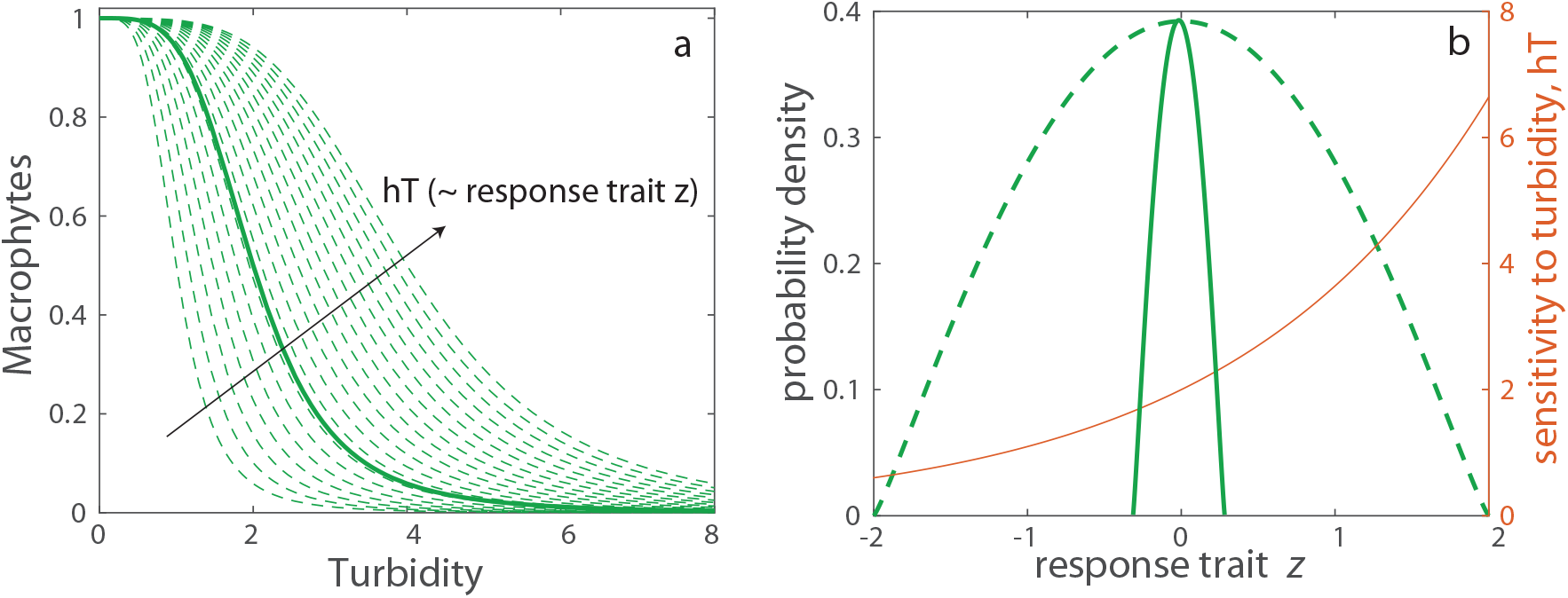
a) Variation in a response trait *z* of macrophytes (e.g. shading tolerance) can affect the way macrophytes respond to water turbidity through parameter *hT* that determines the response of macrophytes to turbidity 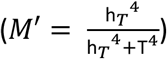. b) Two scenarios of high (dashed) and low (solid) variation in the phenotype distribution of the response trait *z* (~ *Beta*(*μ*, *σ*^2^)), where parameter *h_T_* has a positive relationship with the trait (red line).

